# Impact evaluation of score classes and annotation regions in deep learning-based dairy cow body condition prediction

**DOI:** 10.1101/2022.10.26.513838

**Authors:** Sára Ágnes Nagy, Oz Kilim, István Csabai, György Gábor, Norbert Solymosi

## Abstract

Body condition scoring is a simple method to estimate the energy supply of dairy cattle. Our study aimed to investigate the accuracy with which supervised machine learning, a deep convolutional neural network, can be used to retrieve body condition score (BCS) classes estimated by an expert. Using a simple action camera, we recorded images of animals’ rumps in three large-scale farms. The images were annotated with three different-sized boxes by an expert. A Faster-RCNN pre-trained model was trained on 12 and 3 BCS classes. Training in 12 classes, with a 0 error range, the Cohen’s kappa value yielded minimal agreement. Allowing an error range of 0.25, we obtained a minimum or week agreement. With an error range of 0.5, we had strong or almost perfect agreements. The kappa values of the approach trained on 3 classes show that we can classify all animals into BCS categories with at least moderate agreement. Furthermore, CNNs trained in 3 BCS classes show a remarkably higher proportion of strong agreement than those trained in 12 classes. The prediction precision based on training with various annotation regions showed no meaningful differences.

## 1. Introduction

Body condition scoring of cattle is a widespread, non-invasive, easy-to-use but more or less subjective and time consuming method for estimating an animal’s saturation of the subcutaneous fat stores [1,2]. The different scoring systems infer the animal’s energy supply from the coverage of the lumbar, pelvic, and tail head regions [2]. The saturation of the body stores is quantified using a numerical scale where lean individuals have low values and overweight individuals have high values [3]. The saturation of fat and energy stores can provide important guidance for farm management. Several studies have shown how a shift in body condition score (BCS) away from ideal is associated with changes in production [4]. Too high and low condition points may indicate several abnormalities [4]. The association between a high condition score and the risk of ketosis is well-known [4,5], furthermore, it is also associated with other metabolic problems (e.g. fatty liver) [6], placenta retention [5]. Too low a condition level can be a consequence of lameness [4] and may also be associated with reduced milk production [4]. However, several pathological processes correlate with a decrease in BCS (e.g. metritis, inactive ovaries, displaced abomasum, more days open) [4]. Since pathological changes are often closely related to changes in BCS [4] rather than to a specific condition score, it is understandable that continuous and reliable herd-level condition scoring would be an essential aid to dairy herd management. This is hampered by the fact that scoring requires trained staff [2], herd-level scoring is time-consuming. The volatility of intraobserver and interobserver agreements makes the data generated challenging to use [7].

Our study aimed to investigate the accuracy with which supervised machine learning, specifically deep convolutional neural network (CNN) based Detectron2 models can be used to recover BCS classes estimated by an expert using images of cows taken with a simple RGB camera. As a first approach, we investigated the quality of the predictions of the CNNs taught based on our 12-level BCS scoring for the validation and test set. In the following approach, we investigated the quality of prediction of CNNs trained on three BCS classes corresponding to four different target intervals. Furthermore, we studied how different interest of region rectangles of the rump cause variation in the prediction.

## 2. Materials and Methods

### 2.1. Data collection

Digital video recordings were taken over the course of 2 years with a SJCAM 4000 RGB camera at three large-scale dairy cattle farms in Hungary (farm F1: 1150 cows, F2: 880 cows, F3: 960 cows) with the camera positioned in the rotary milking parlour pointing at the animals’ rump. Not all of the milking cows were filmed in each video. To avoid over-representation of any cow with a given body scoring in the data set, we skipped at least a month between any two videos taken at a single site to ensure the conditions of the animals had changed.

### 2.2. Data pre-processing

The videos were annotated by an expert remotely using the Visual Object Tagging Tool (VoTT, v2.2.0) [8], assigning both a bounding rectangle to the animals rump as well as an estimated BCS. Scoring was performed in the range of 1-5 BCS. In this range 12 levels were defined: 1 to 2.5 and 4 to 5 were divided into 0.5 score intervals while between 2.5 and 4.00 were divided more finely with 0.25 score intervals [2]. The scoring was done continuously with movie footage, drawing the bounding boxes and making the BCS inference from only the image captured where the rump was at the closest point to the camera, these were the final images used to build the train, validation and test sets. After annotating the videos produced in the study, the same expert rechecked and adjusted the annotations by Label Studio [9], where all images were managed together. Farm F1 and F2 images were split into the train and validation sets. This was done with stratification of BCS scores; randomly selecting 80% of the images within each score level for the train set, with the remaining pictures, we created the validation set. This ensured the training and validation sets contained the same class distributions so the validation set loss would be appropriate tool for model choice decisions. The annotated images from site F3 were retained as an independent test set. The number of images per score in each set is summarized in Table 1. Three sizes of bounding box were generated for each image automatically from initial annotations (See supplementary material: Automated 3 box size annotation).

**Table 1.** The number of annotated images included in the study per score. Images from sites F1 and F2 were used to create the training and validation set, while the held out test set consisted of images from the F3 site only.

| BCS | F1 & F2 farm<br>train | validation | F3 farm<br>test |
| --- | --- | --- | --- |
| 1.00 | 162 | 41 | 22 |
| 1.50 | 215 | 54 | 31 |
| 2.00 | 363 | 91 | 53 |
| 2.50 | 370 | 93 | 65 |
| 2.75 | 182 | 46 | 42 |
| 3.00 | 323 | 81 | 41 |
| 3.25 | 381 | 95 | 50 |
| 3.50 | 659 | 165 | 66 |
| 3.75 | 74 | 18 | 9 |
| 4.00 | 58 | 14 | 11 |
| 4.50 | 46 | 12 | 6 |
| 5.00 | 89 | 22 | 11 |
| Total: | 2922 | 732 | 407 |

### 2.3. Choice of model architecture

From machine learning’s viewpoint the body scoring is an object detection and classification problem. For this reason we chose an object detection model that can leverage this attention focus in the form of both bounding boxes and classes. The Faster-RCNN architecture [10] has a shared model internal representation for localisation and classification prediction joint tasks which matched exactly our problem setting. This joint learning task is apparent by inspecting the form of the loss function used for training *ℓ* = *ℓ*_*cls*_ + *ℓ*_*bbox*_. Due to state of the art results, the Detectron2 [11] implementation of the Faster-RCNN architecture was chosen. All models were downloaded from the Model Zoo code repository with network parameters pre-trained on COCO Dataset.

### 2.4. Evaluation metrics

The performance of our model’s localisation can be reviewed in terms of bounding box prediction average precision at IoU=0.50 (AP50). This value increases when predicted bounding boxes overlap more with ground truth annotation bounding boxes. In addition to detecting an object in the image, Detectron2 estimates class probability distribution over all classes. As final classification it assigns the object to the class for which the probability is the highest among all possible classes. The quality of the predictions was quantified using Cohen’s kappa [12] and accuracy. The value of *kappa* = (*P*_0_ *− P*_*e*_)/(1 *− P*_*e*_) where *P*_0_ is the observed agreement between ground truth and predicted classes and *P*_*e*_ is the probability change agreement between model prediction and annotation ground truth. Cohen’s kappa values can be interpreted: 0 0. *−* 20 no, 0.21 *−* 0.39 minimal, 0.40 *−* 0.59 weak, 0.60 *−* 0.79 moderate, 0.80 *−* 0.90 strong, above 0.90 almost perfect agreement [13]. Following the “one-versus-all” accuracy was calculated comparing each class to the remaining levels by the formula *accuracy* = (*TP* + *TN*)/(*TP* + *TN* + *FP* + *FN*), and from those an overall accuracy reported as the mean of them.

### 2.5. Model screening

In order to pre-screen for the most appropriate pre-trained model for the BCS task, 10 pre-trained models of Detectron2 [11] were further trained and validated on our data sets each with identical hyperparameters for 15 epochs. In this model selection phase, raw BCS scores were to be classified into three classes (BCS classes: <2.5, between 2.5 and 3.75, >3.75). The `faster_rcnn_R_50_FPN_3x` model gave the lowest validation loss, so we selected and used this pre-trained model in all further experiments (Table 2).

**Table 2.** Model selection. Ten pre-trained models of Detectron2 were run on the same data set with the same settings (number of epochs: 15). The model with the lowest validation loss was chosen for all further experiments.

| Pretrained model | Validation loss |
| --- | --- |
| faster_rcnn_R_50_FPN_3x | 0.0612 |
| faster_rcnn_R_101_FPN_3x | 0.0628 |
| faster_rcnn_R_50_FPN_1x | 0.0637 |
| faster_rcnn_X_101_32x8d_FPN_3x | 0.0662 |
| faster_rcnn_R_50_DC5_1x | 0.0796 |
| faster_rcnn_R_50_DC5_3x | 0.0840 |
| faster_rcnn_R_101_C4_3x | 0.0848 |
| faster_rcnn_R_101_DC5_3x | 0.0848 |
| faster_rcnn_R_50_C4_1x | 0.1019 |
| faster_rcnn_R_50_C4_3x | 0.1040 |

### 2.6. Model training and prediction

#### 2.6.1. With 12 BCS classes

Using the selected pre-trained model the first experiment was to run by training validation and testing on the 12-level ordinal scoring annotations. This was repeated for each of the three square sizes (see Fig.1) separately. Using the model’s optimal weights at checkpoints with the lowest validation losses and AP50s we made predictions on the validation and test sets. The value where the probability distribution (output by the model) had it’s maximum was taken as the estimated body score. We also kept the “predicted class probabilities” as they are a measure of model confidence for a given prediction.

Following the approach of Yukun et al. (2019) [14], we evaluated the model predictions with 0, 0.25 and 0.5 error thresholds to account for the ordinal nature of the body condition scores and thereby allowing “near miss” predictions to be defined as correct, reducing metric stringency.

**Figure 1.**
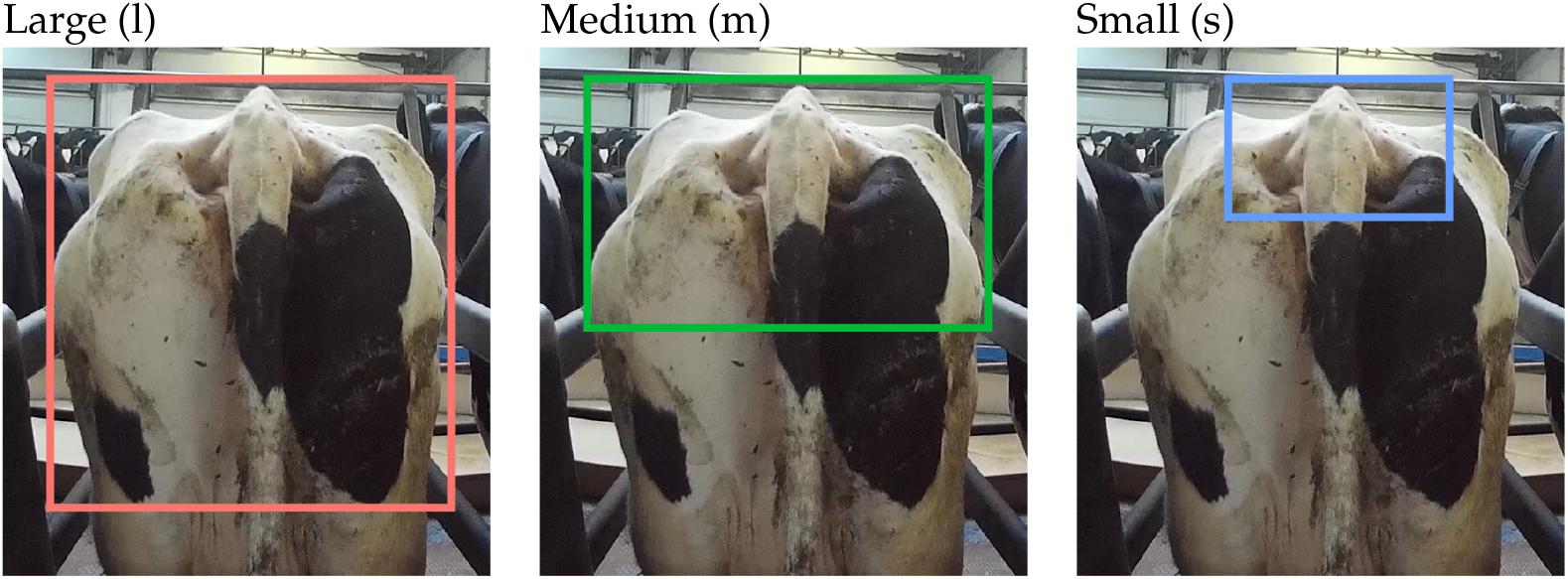
Annotation rectangles. The large (l) box in the image of the animal was placed across the entire width of the rump from the tail head to mid-thigh. The medium (m) one was the entire width of the rump from the tail head to the base of the vulva. While the small (s) box frames the ischial tuberosities and the tail head only.

#### 2.6.2. With 3 BCS classes for 4 practical target intervals

In addition to the 12-point BCS annotation, we also assessed the quality of the predictions according to more broad BCS classes of practical relevance. Different BCS ranges are considered optimal at different stages of lactation. These target intervals are summarized in Table 3. We re-labelled all the original BCSs according to these 4 different threshold regimes (T1,T2,T3,T4) target ranges. T1 optimal target interval for calving (days in milk (DIM): 0), dry (DIM: -60 to -1) and dry off (DIM: >300); T2 for early (DIM: 1 to 30) and mid (DIM: 101 to 200) lactation; T3 for peak lactation (DIM: 31 to 100) while T4 for late lactation (DIM: 201 to 300). We created three new BCS classes for each type of thresholdings: below target interval, target interval and above target interval. The training, validating and testing sessions were re-run with these new 3 classes for each of the 4 thresholding regimes and for each of the 3 bounding box sizes. Each experiment used the same input images just the label definitions varied between each. In Table 3 the linked figure shows the proportion of images after reclassification in the train, validation and test sets. The proportion of classes is the same in the training+validation set (farm F1+F2) and test (farm F3) set. This allows for a fair testing as the class imbalance is the same in train+validation and test sets.

**Table 3.**
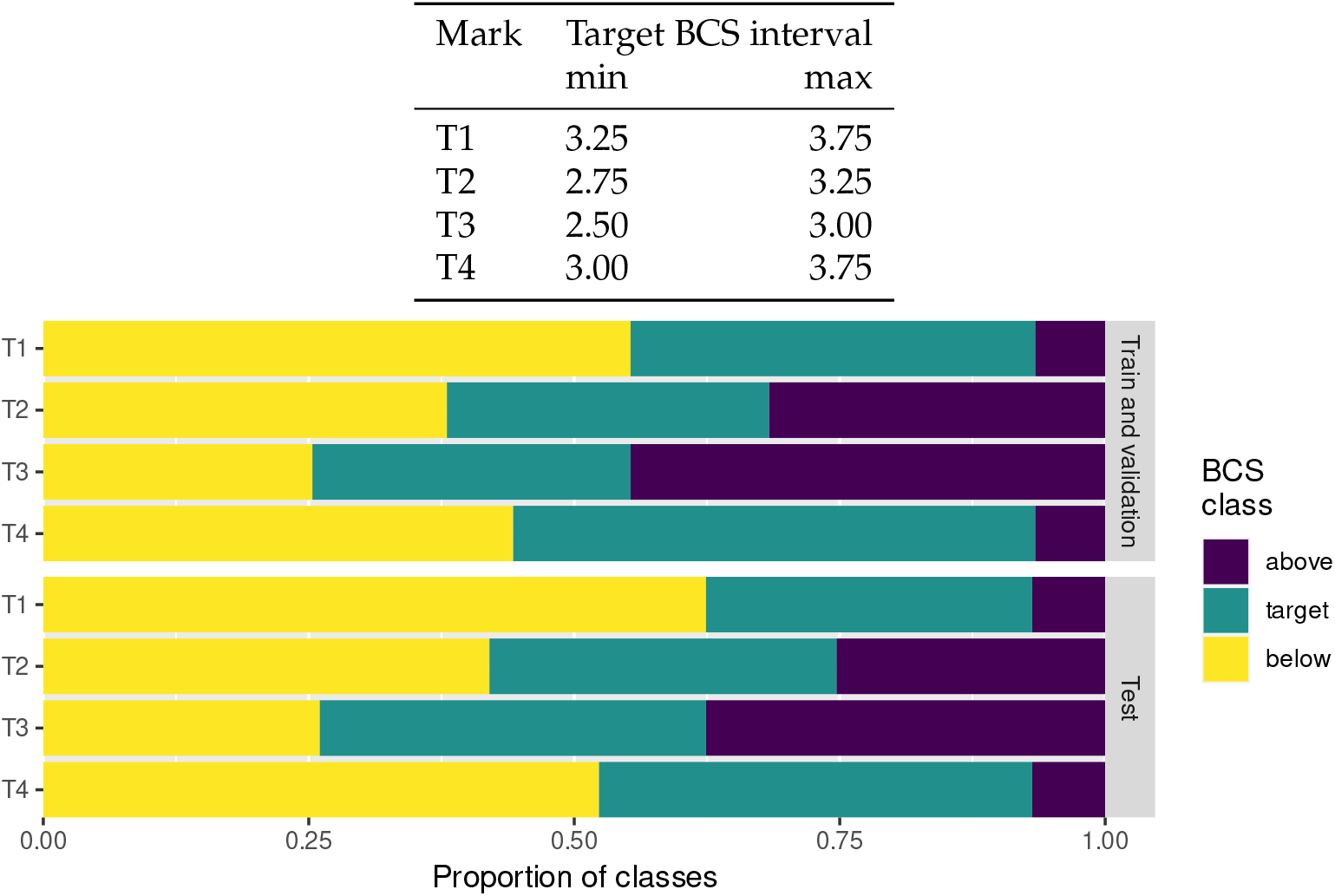
Body condition score target intervals. T1 for calving (DIM: 0), dry (DIM: -60 to -1) and dry off (DIM: >300). T2 for early (DIM: 1 to 30) and mid (DIM: 101 to 200) lactation. T3 for peak lactation (DIM: 31 to 100) and T4 for late lactation (DIM: 201 to 300). Below shows the re-labeling for each interval. For each regime the original ordinal labels are re-labeled according to the given threshold of that regime. For example: With the T1 thesholding there are more images with cows in the “below” class whereas if we re-label the data with the T2 thesholding then the 3 classes are more evenly split. Class distributions in training/validation sets match respective test sets.

To compare the training process between training on 3 classes and training on 12 classes we repeated the analysis in a way where 12 classes were predicted but they were then re-classified to the new 3 classes at inference time (see Figure 6).

## 3. Results

### 3.1. Prediction with 12 BCS classes

The quality of the predictions for the 12 classes, based on the training with 12-classes, is shown in Figure 2. The *x* axis represents the results thresholded by predicted class probabilities. As the threshold increases only images classified with high model confidence are used to calculate kappa and accuracy. Traversing the *x* axis of each plot gives an idea of how the data is clustered near the learnt decision boundary in high dimensional space; images near to the decision boundary have a low predicted class probability (model confidence). Images with high confidence are to be classified better.

**Figure 2.**
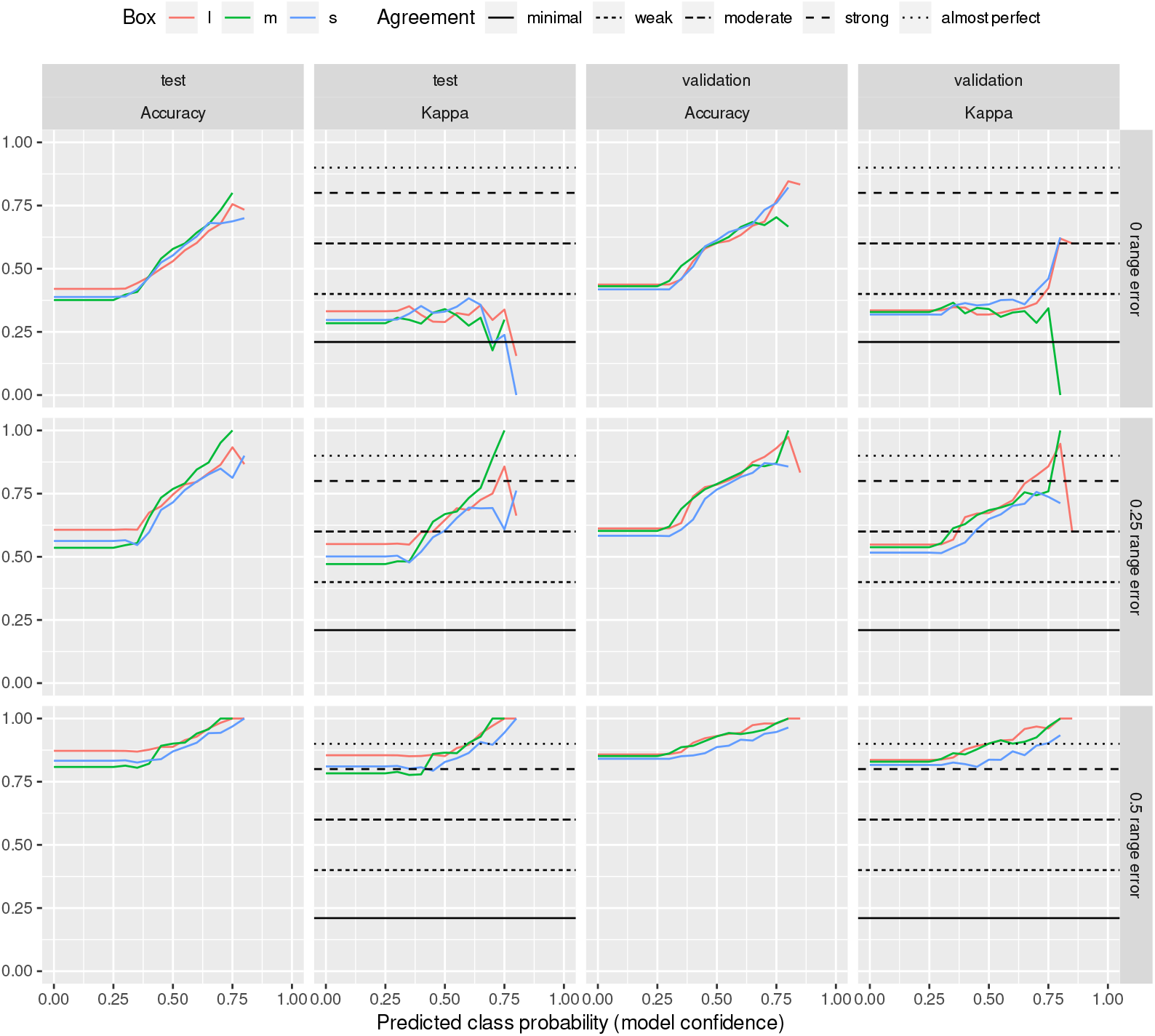
Model trained and evaluated with 12 BCS classes. For the agreement analysis on the expert given and predicted scores three error ranges (0, 0.25 and 0.5) were alowed. The horizontal lines represent the thresholds of McHugh[13] to interpret Cohen’s kappa values.

Training and evaluating in 12 classes, allowing for an error range of 0, the kappa value on the test set yielded a minimal agreement but worse agreement above a class prediction probability of about 75%. For the validation set, the agreement is similar to below 75% class prediction probability. In contrast, above this level, the agreement is weak for boxes l and s and even weaker than the minimum for box m. If we allowed an error range of 0.25, we obtained a minimum agreement below the 50% class prediction probability on both the test and the validation set, and above that, the maps fall into the weak agreement range. Allowing an error range of 0.5, we have curves running above or close to the strong cut-point with a class prediction probability of about 60% on both the test and validation sets. However, from 65-70%, we get almost perfect agreement.

### 3.2. Prediction with 3 BCS classes

Figure 3 shows the quality of the predictions made on only 3 BCS classes instead of 12 BCS classes. The three classes were created according to the four target ranges (T1-T4) presented in Table 3. The expert’s scores assigned to the images were reassigned to one of the three BCS classes and then used to perform the training, with the prediction also being made for three classes. In this 3 class regime the network makes high quality predictions. This type of classifier may be more relevant for farms to give lower resolution but reliable predictions and act more as a screening tool. If animals are predicted as out of target they could be further investigated.

**Figure 3.**
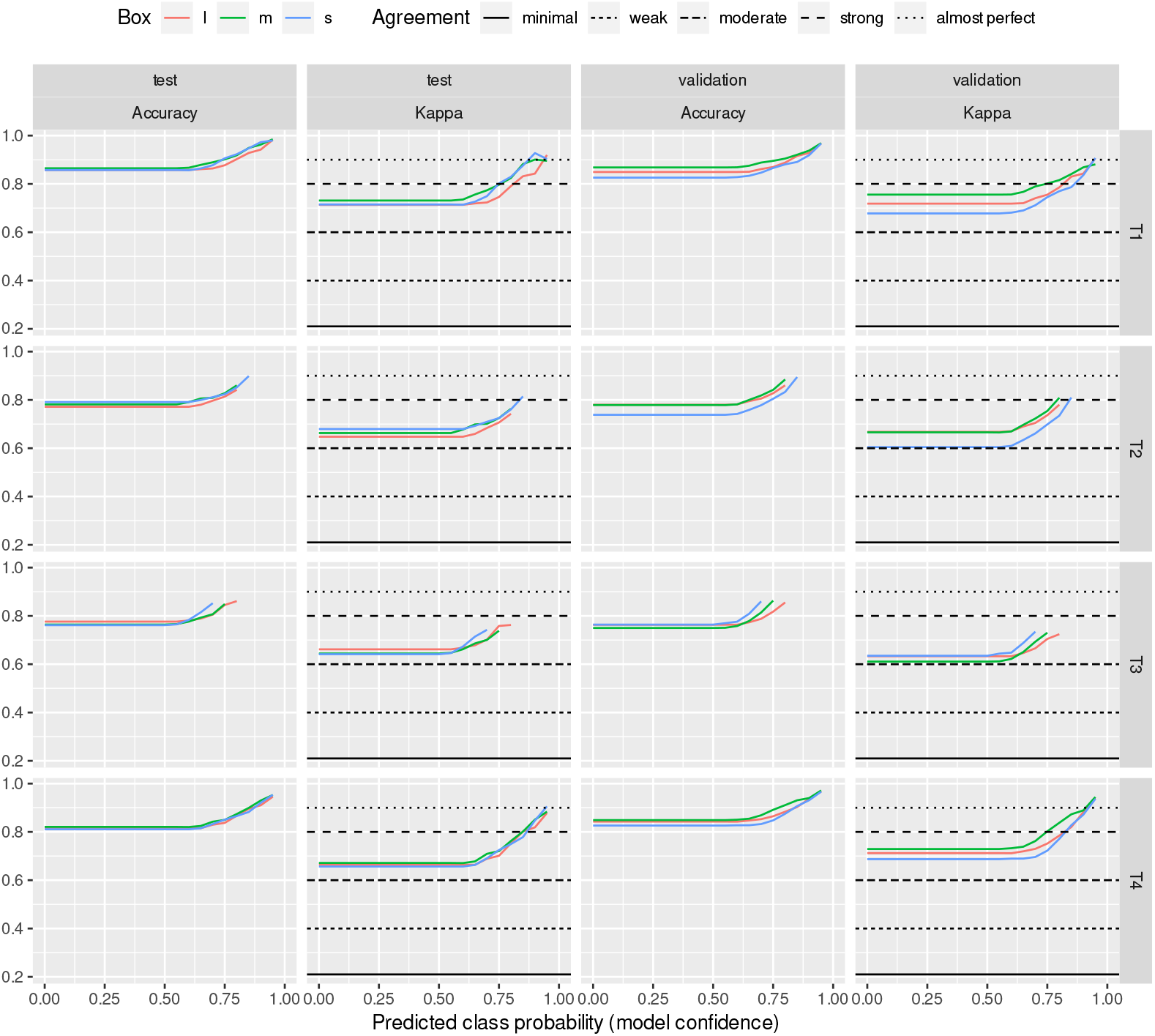
Model trained and evaluated with 3 BCS classes. The horizontal lines represent the thresholds of McHugh[13] to interpret Cohen’s kappa values.

In a second “control” approach we trained the network on 12 BCS classes and returned 12 classes with the predictors as in Figure 2. We then re-classified the 12 classes into the three BCS classes. This was also repeated for each T1-4 regime. These results can be seen in Figure 6. The traversal of model confidence thresholding shows more noise. This is understandable as the body scoring classes are ordinal but not continuous. To further compare the two approaches, we examined the difference in kappa and accuracy values for the joint class classification probabilities. For that we subtracted the values of trained on 12 classes from the values of trained on 3 classes. In Figure 7/a the mean and the standard deviation of differences are plotted this is outlined in the supplementary material section. The methods show similar performances.

The differences between the prediction goodness of annotation boxes obtained from the networks taught by the 3 classes are summarized in Figure 7/b. We may observe the same trends for accuracy and kappa for all target ranges in box prediction differences. It can also be seen that the differences between the boxes are an order of magnitude smaller than those obtained from the networks taught by classes 12 and 3. Based on the test set, the prediction precision per box in descending order for the target ranges is as follows: T1: m, s, l; T2: s, m, l; T3: l, s, m; T4: m, l, s. In the validation set T1: m, l, s; T2: m, l, s; T3: s, l, m; T4: m, l, s.

As we traverse the *x* axis for each result, we have fewer and fewer predicted images to evaluate the metric. Figure 4 summarises the proportion of the initial (test or validation) data set corresponding to each predicted class probability value. For the neural networks trained on 12 classes, already at a predicted class probability value of 40%, half of the images are dropped. However, for networks taught by 3 classes, even at the highest predicted class probability value, half of the images or slightly less than half of the images are still part of the analysis. The 3 class classification can be thought of as a more “easy” task for the network and so higher model confidence for the predicted classes is reasonable. We find variation between results of the 3 box sizes not to be very large indicating that the supervised signal is mainly found within the medium box area.

**Figure 4.**
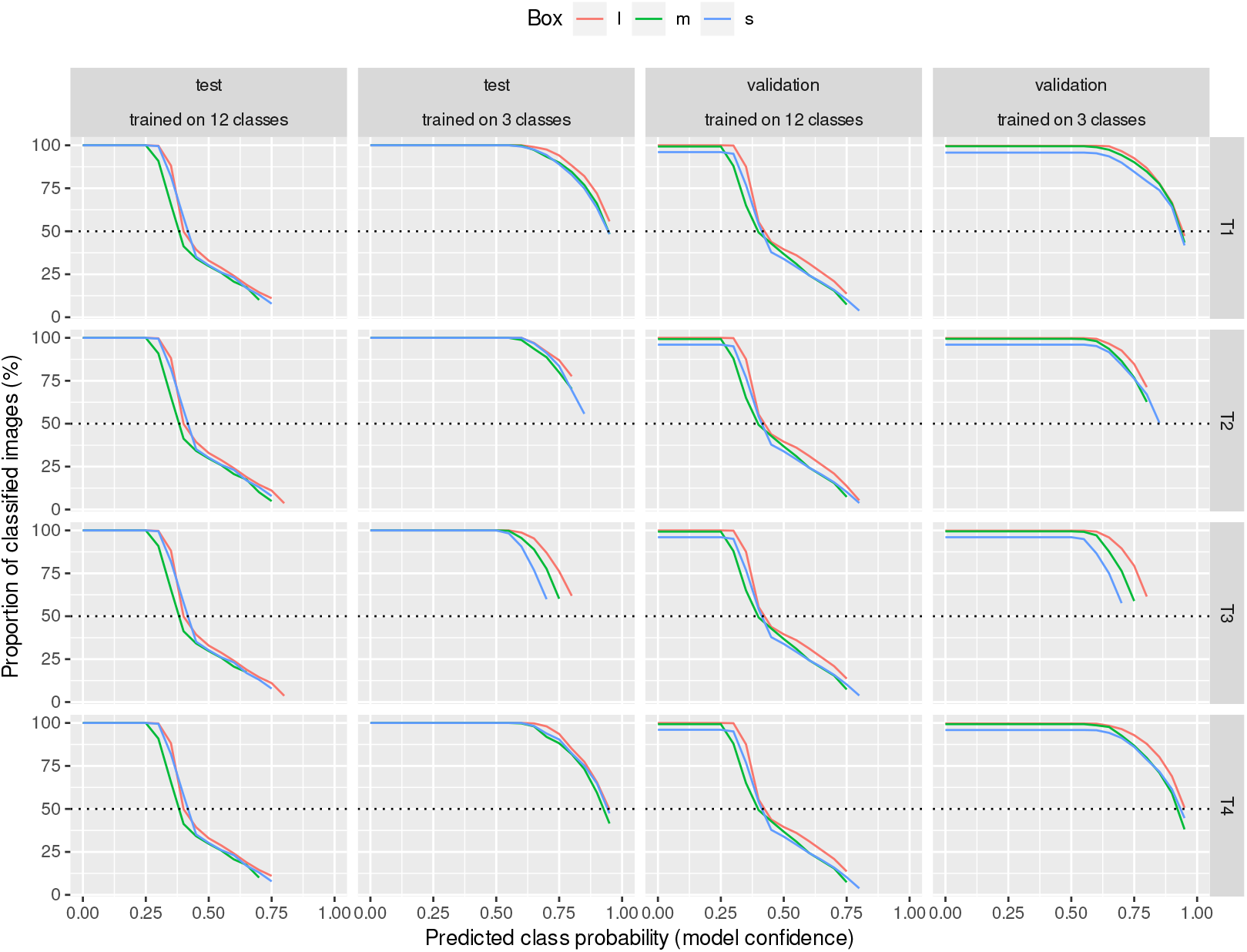
Proportion of classified images. As threshold cutoff is more stringent, the number of images left gets smaller. The reduction occurs to a larger extent for the 12 class experiments. This gives an insight into the distribution of images with respect to the learnt decision boundary in the model representation.

Regardless of the target range (T1-T4), the kappa values of the approach trained and evaluated in the 3 classes show that we can classify all animals into BCS categories with at least moderate agreement. The Figure 5 shows the proportion of predictions based on training for different target ranges with different box sizes that resulted in at least strong agreement.

**Figure 5.**
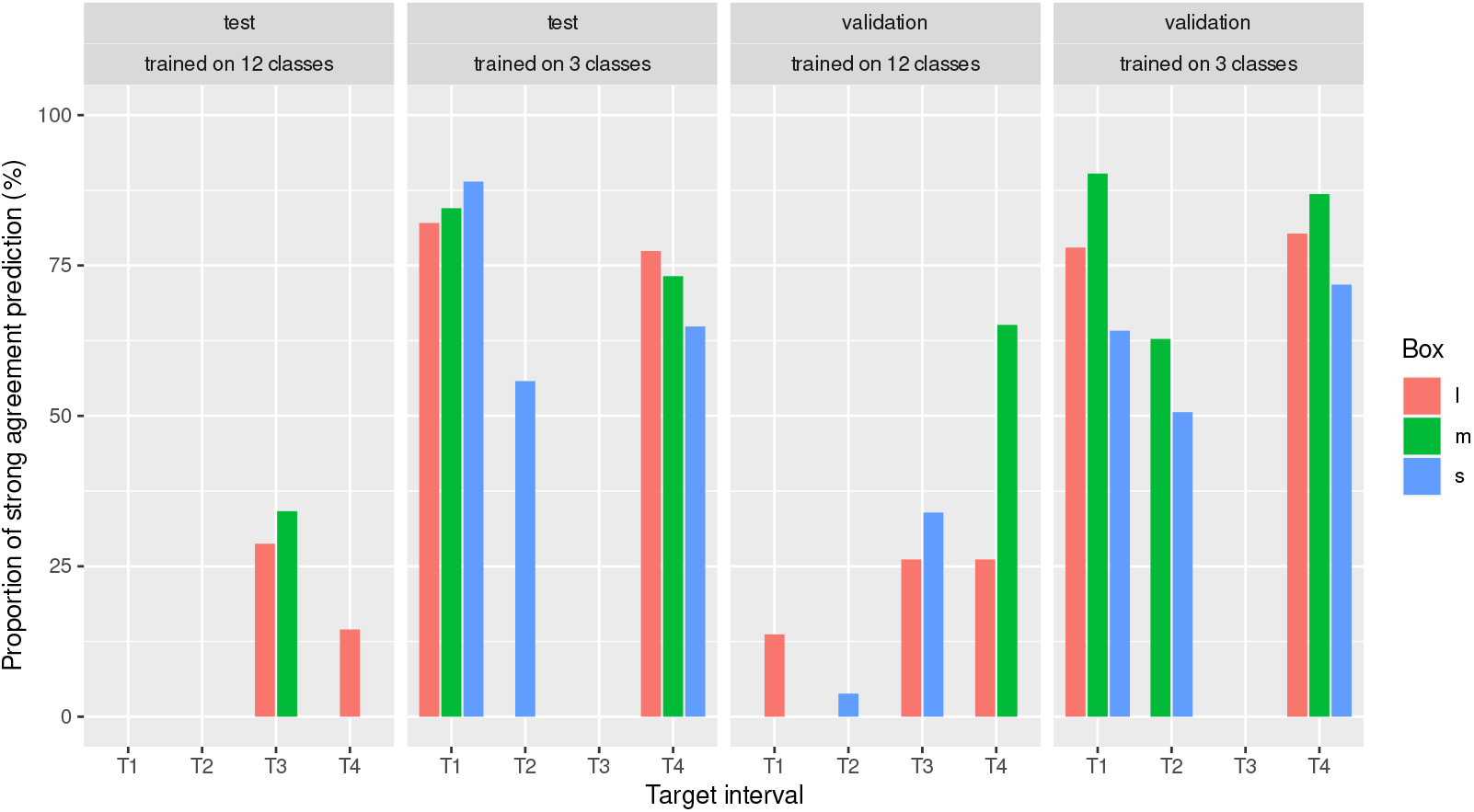
Proportion of predictions with strong agreement (Cohen’s kappa *≥* 0.8). For predictions over different target ranges (T1-T4), CNNs trained in 3 BCS classes show a remarkably higher proportion of strong agreement than those trained in 12 classes and then reclassified.

**Figure 6.**
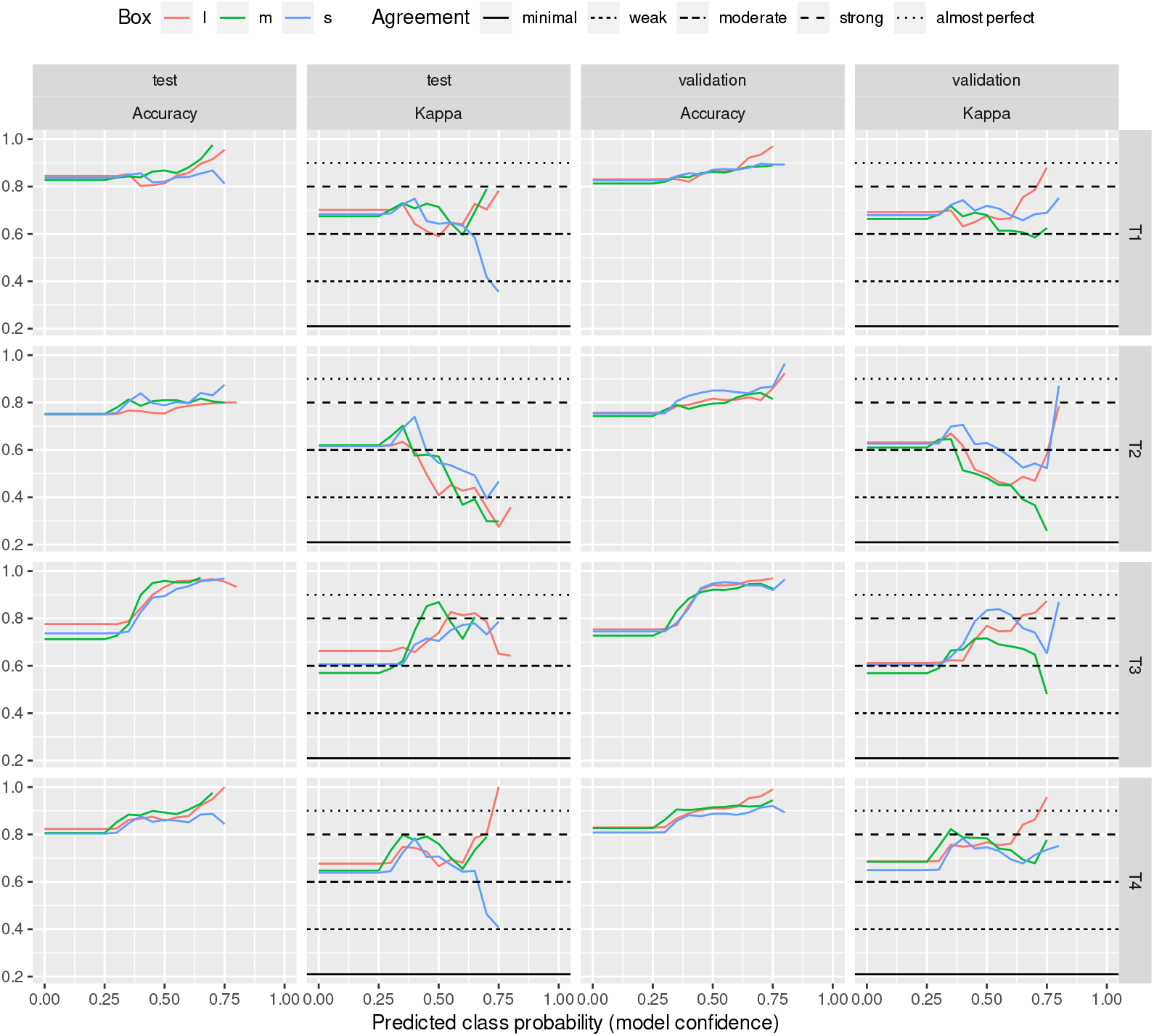
Model trained and evaluated with 12 and 3 BCS classes, respectively. The horizontal lines represent the thresholds of McHugh[13] to interpret Cohen’s kappa values.

**Figure 7.**
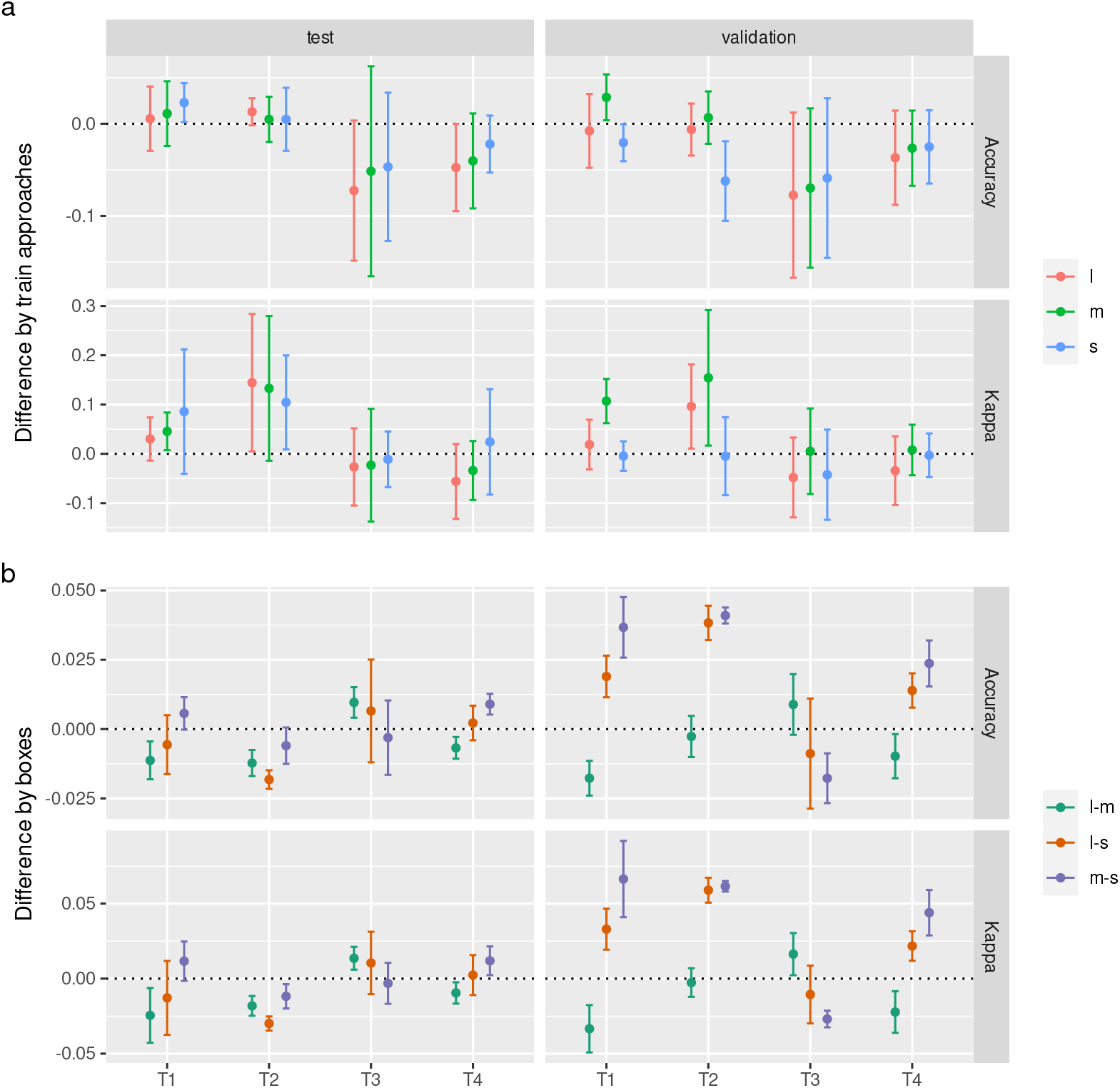
Prediction precision variability shown by mean and standard deviation. Sub-figure a represents the difference between the predictions based on the 3- and 12-class training. Differences among the box sizes in prediction precision obtained from the 3-class train are summarized in sub-figure b.

## 4. Discussion

Several possible approaches to estimating cattle body conditions using neural networks exist [14–16]. The more common approach in the literature is to use recordings of animals scored with high-resolution scoring, for example, 0.25 or 0.5-unit increments, to train the neural network and to test how the trained network performs in terms of prediction reliability at the same scale. Assume no discrepancy is allowed between observed and predicted scores. In that case, these approaches yield a weak agreement, as indicated by the *kappa* = 0.45 value of Yukun et al.[14] If we allow some variation between observed and predicted scores, both kappa and accuracy values improve. As the first step in our investigation, we followed the approach, and similar results were obtained around the reliability values presented by other authors.[14,15,17–21]

However, in addition to these high-resolution score classes, we felt it worthwhile to investigate the prediction quality that can be achieved for practically important condition score classes. We conducted this investigation in two forms. In one, we trained a neural network on the high-resolution, detailed twelve-point level dataset, and then both predictions and original expert scores were classified into three condition categories. Thus, there were a target range category (T1, T2, T3, T4) and categories below and above the target range. In the second approach, we trained the neural network with the data set already divided into the three categories and made predictions corresponding to the three classes. The test and the validation set’s predictions show that the latter approach gave better results with lower noise. However, after splitting into three categories, the networks trained on the 12 categories also gave good results, but these results were noisier and can be evaluated as a less robust approach.

We also considered the probability that the neural network assigned a BCS class to the object detected in each image. The kappa and accuracy values presented were evaluated as a function of this class assignment probability. It can be seen that the precision of the predictions improves with increasing class assignment probability. Nevertheless, as the class ranking probability increases, fewer images can be considered in evaluating the reliability of the predictions. Comparing networks trained on 12 classes and 3 classes, it can be seen that the class assignment probabilities are lower for the former than for the latter. For this reason, as we increase the threshold of the classification probability of the images included in the prediction precision analysis, the number of usable images predicted by the neural network trained on the 12 classes decreases rapidly. While for networks trained on 3 classes, the number of usable images decreases much more slowly as the classification probability threshold increases. Even at the highest threshold, roughly half of the images were retained. This approach has not been found in the literature where the prediction precision is analyzed in conjunction with the classification probability. However, the results show that it significantly affects the prediction quality and, thus, the practical efficacy of body condition prediction based on neural networks. Nevertheless, it is still problematic that the number of images with a higher classification probability is less than the number of animals in the set used for prediction. The results show that at the most reliable classification probability threshold, we lose half of the animals, which means we obtain reliable conditioning information for half of the animals on a given day. However, we aim to obtain daily information on all individuals in a given herd. We see several possibilities to address this problem, which further studies could clarify. One approach could be based on the fact that animals are snapped not only once but several times during milking in the carousel systems. We could identify the highest probability class from their class distribution if we predicted each of these. Nevertheless, we can conclude from Figure 5 that the prediction of CNNs trained in class 3 shows a significant proportion of strong agreement, which is better than the inter-rater agreement found in the literature.

The use of practical thresholds in the three-grade approach trained in 12 classes and assessed in three classes is problematic. Several authors show that high prediction reliability can be obtained in error ranges of 0.25 or 0.5.[14,15,17–21] However, when we try to assign this to the practical condition categories we use, it is impossible for many animals to decide which practical condition interval to place an animal in with an error range of 0.25 or 0.5. It is important to emphasize that our study aimed to investigate the reliability with which a neural network can reproduce the scores and score categories of animals scored by an expert. It is also possible to construct a ground of truth from the scores of several experts rather than one. However, it is worth considering that the agreement between the scores of two independent experts is weak or moderated based on the literature. The results reported by Mullis et al.[22] show that the Cohen’s kappa of the agreement between two experts’ BCS values is 0.62 and 0.66, while Song et al.[23] found that an inter-assessor agreement kappa=0.48, while the intra-assessor agreement kappa is 0.52 and 0.72. Thus, the prediction precision of a neural network built on this basis could easily be worse, not better.

In our study, we used a test set from an independent site in addition to the validation set to see the robustness of the neural network prediction. After all, it is expected that if animals from the same farm are given the training and validation sets, the prediction for the validation set will be better than the predictions for an utterly independent farm. Surprisingly, the predictions of the networks trained on the three classes differ little for the test and validation sets. A further interesting feature of our results is that the prediction precision based on the three types of annotation boxes (large, medium and small) also differ very little. Our reasoning in choosing three different-size annotation boxes was that we might think that a large box contains more information that the algorithm can capture. Conversely, it seems that a medium-sized box contains the most usable information, which still has little noise. Next in line is the small box, which gives slightly better results than the large box. Here we can think of it as containing less information and less noise. In contrast, the big one has more noise with more information. Thus, the order is that the middle one comes after the small and then the large box in prediction performance.

In our work, we deliberately did not use complementary tactile examinations in the generation of expert scores because when teaching a neural network based purely on images, this information is not available to the algorithm, so the expert’s information in scoring is richer than what we can offer the neural network.

Our results show that the quality of training and prediction from two-dimensional images taken with a simple sports camera using Detectron2 is not inferior to prediction results based on three-dimensional cameras or on scoring with tactile detection. An additional option to consider to improve the prediction quality could be to use ensemble prediction of different trained networks as the final output.

Our work resulted in weights generated for the neural network and made publicly available. Others can use this as a pre-trained network in training neural networks on similar images. Thus, presumably, they can create their neural network using fewer images to predict BCS categories.

## 5. Conclusion

Our results conclude that CNN training on classes corresponding to practically relevant target ranges gives more robust and precise predictions than training on highresolution classes. With predictions based on target interval training, we obtained similar or even better results than the agreement between experts. The prediction precision based on training with various annotation regions showed no meaningful differences.

## Author Contributions

NS takes responsibility for the integrity of the data and the accuracy of the data analysis. NS, SÁG and GG conceived the concept of the study. SÁG performed the data annotation. NS and SÁG participated in the model training, predictions and statistical analysis. NS, SÁG and OK participated in the drafting of the manuscript. NS, SÁG, OK, IC and GG carried out the critical revision of the manuscript for important intellectual content. All authors read and approved the final manuscript.

## Institutional Review Board Statement

Not applicable.

## Informed Consent Statement

Not applicable.

## Data Availability Statement

The weights used for the predictions from training with each BCS class and annotation box combination can be downloaded from: https://doi.org/10.6084/m9.figshare.21372000.v1

## Acknowledgments

We thank Alex Olár for his suggestions to help us in our work.

## 6. Supplementary material

### 6.1. Automated 3 box size annotation

After the initial annotation, the position and size of the rectangles on the images were standardized as follows. Three rectangles of different sizes were placed on the images of the F3 farm by the expert annotator (Fig 1). The large (l) box in the image of the animal was placed across the entire width of the rump from the tail head to mid-thigh. The medium (m) box contained the entire width of the rump from the tail head to the base of the vulva. While the small (s) box frames the ischial tuberosities and the tail head. We created a training set from 80% of these images and 20% a validation set. These were used to train the `faster_rcnn_R_50_FPN_3x` pre-trained model of the Detrectron2 environment model zoo. During the training, we recorded the validation loss and AP values every 100 iterations and kept the weights that gave the best results. Using the trained model, we predicted the three types of annotation squares for all images and used them for all the main experiments. These bounding boxes were validated visually by the expert annotator.

### 6.2. 12 class training and testing after re-labeling test sets into T1-4 classes

Within the test set, for both the T1 and T2 target ranges, the kappa and the accuracy are slightly higher for networks taught in 3 classes than for those taught by 12 classes. For T3 and T4, the opposite is seen. The kappa shows the same trend, with the difference that for the target range T4, the s box shows a higher value than the network taught by 3 class. Within the validation set, the accuracy values of the predictions taught by 12 classes were generally higher. The exception is the m box for the T1 and T2 target domains. For all target domains, the s box resulted in higher kappa for the networks taught by 12 classes. In the target ranges T1 and T2, the l box resulted in a higher kappa for networks taught in 3 classes, while it resulted in a lower kappa for the target ranges T3 and T4. The kappa values of the m boxes were higher for CNNs taught by 3 classes in all target ranges.

